# Electrically Conductive Pili from Pilin Genes of Phylogenetically Diverse Microorganisms

**DOI:** 10.1101/118059

**Authors:** David J.F. Walker, Ramesh Y. Adhikari, Dawn E. Holmes, Joy E. Ward, Trevor L. Woodard, Kelly P. Nevin, Derek R. Lovley

**Affiliations:** Department of Microbiology; Department of Physics University of Massachusetts, Amherst, MA 01003; Department of Physical and Biological Sciences, Western New England University, Springfield, MA

**Keywords:** Geobacter, Flexistipes, Calditerrivibrio, Desulfurivibrio, Desulfofervidus, e-pili

## Abstract

The possibility that bacteria other than *Geobacter* species might contain genes for electrically conductive pili (e-pili) was investigated by heterologously expressing pilin genes of interest in *Geobacter sulfurreducens*. Strains of *G. sulfurreducens* producing high current densities, which are only possible with e-pili, were obtained with pilin genes from *Flexistipes sinusarabici, Calditerrivibrio nitroreducens*, and *Desulfurivibrio alkaliphilus*. The conductance of pili from these strains was comparable to native *G. sulfurreducens* e-pili. The e-pili derived from *C. nitroreducens,* and *D. alkaliphilus* pilin genes are the first examples of relatively long (> 100 amino acids) pilin monomers assembling into e-pili. The pilin gene from *Desulfofervidus auxilii* did not yield e-pili, suggesting that the hypothesis that this sulfate reducer wires itself to ANME-1 microbes with e-pili to promote anaerobic methane oxidation should be reevaluated. A high density of aromatic amino acids and a lack of substantial aromatic-free gaps along the length of long pilins may be important characteristics leading to e-pili. This study demonstrates a simple method to screen pilin genes from difficult-to-culture microorganisms for their potential to yield e-pili; reveals new potential sources for biologically based electronic materials; and suggests that a wide phylogenetic diversity of microorganisms may employ e-pili for extracellular electron exchange.

## Introduction

Electrically conductive pili (e-pili) can play an important role in the biogeochemical cycling of carbon and metals and have potential applications as ‘green’ electronic materials (Lovley 2011, Lovley and Malvankar 2015, Lovley 2017, Malvankar and Lovley 2014, Shi et al 2016). Therefore, a better understanding of the diversity of microorganisms capable of producing e-pili has many potential benefits. To date, e-pili have been most intensively studied in *Geobacter sulfurreducens* and the closely related *Geobacter metallireducens* (Adhikari et al 2016, Feliciano et al 2015, Lampa-Pastirk et al 2016, Malvankar et al 2011, Malvankar et al 2014, Malvankar et al 2015, Reguera et al 2005, Tan et al 2016b, Tan et al 2017, Vargas et al 2013, Xiao et al 2016). These studies have demonstrated that the e-pili of both *G. sulfurreducens* and *G. metallireducens* are sufficiently conductive along their length, under physiologically relevant conditions, to account for maximum potential rates of extracellular electron transfer (Adhikari et al 2016, Lampa-Pastirk et al 2016, Tan et al 2016b, Tan et al 2017). *Geobacter* strains that lack e-pili, or express pili with diminished conductivities, are incapable of direct interspecies electron transfer (DIET) and Fe(III) oxide reduction (Liu et al 2014, Reguera et al 2005, Rotaru et al 2014a, Rotaru et al 2014b, Shrestha et al 2013, Summers et al 2010, Tan et al 2016b, Tremblay et al 2012, Vargas et al 2013). e-Pili are also required for *Geobacter* species to generate high current densities when electrodes serve as the sole electron acceptor (Liu et al 2014, Reguera et al 2006, Tan et al 2016b, Vargas et al 2013). Increasing e-pili expression increases the conductivity of electrode biofilms and their current output (Leang et al 2013, Yi et al 2009).

Aromatic amino acids are key components in electron transfer along the length of *G. sulfurreducens* e-pili (Adhikari et al 2016, Lampa-Pastirk et al 2016, Tan et al 2016a, Tan et al 2017, Vargas et al 2013). There is debate whether the aromatic amino acids contribute to a traditional electron-hopping electron transport mechanism or whether over-lapping π-π orbitals of the aromatics confer a metallic-like conductivity similar to that observed in synthetic conducting polymers (Lampa-Pastirk et al 2016, Lovley and Malvankar 2015, Malvankar et al 2011, Malvankar and Lovley 2014, Malvankar et al 2014, Malvankar et al 2015, Vargas et al 2013). However, the uncertainty over these mechanistic details should not obscure the fact that long-range electron transport via e-pili is a remarkable strategy for long-range biological electron transfer.

It has been speculated that the relatively small size (61 amino acids) of the *G. sulfurreducens* pilin is a key feature contributing to its assembly into e-pili (Holmes et al 2016, Malvankar et al 2015, Reguera et al 2005). Pili comprised of larger pilins, such as those found in *S. oneidensis* (Reguera et al 2005), *Pseudomonas aeruginosa* (Liu et al 2014, Reguera et al 2005), and *G. uraniireducens* (Tan et al 2016b) have poor conductivity. Based on these results, it was proposed that smaller pilins might permit the tight packing of aromatic amino acids required for conductivity (Malvankar et al 2015, Xiao et al 2016). However, previously investigated poorly conductive pili comprised of longer pilins have either: 1) a lower overall density of aromatic amino acids than those found in e-pili (i.e. *P. aeruginosa* and *S. oneidensis)* or 2) a large gap within the pilin sequence devoid of aromatic amino acids, such as observed in the pilin of *G. uraniireducens* (Figure 1). These considerations suggest that the conductivity of a broader diversity of pili should be investigated.

**Figure 1.**
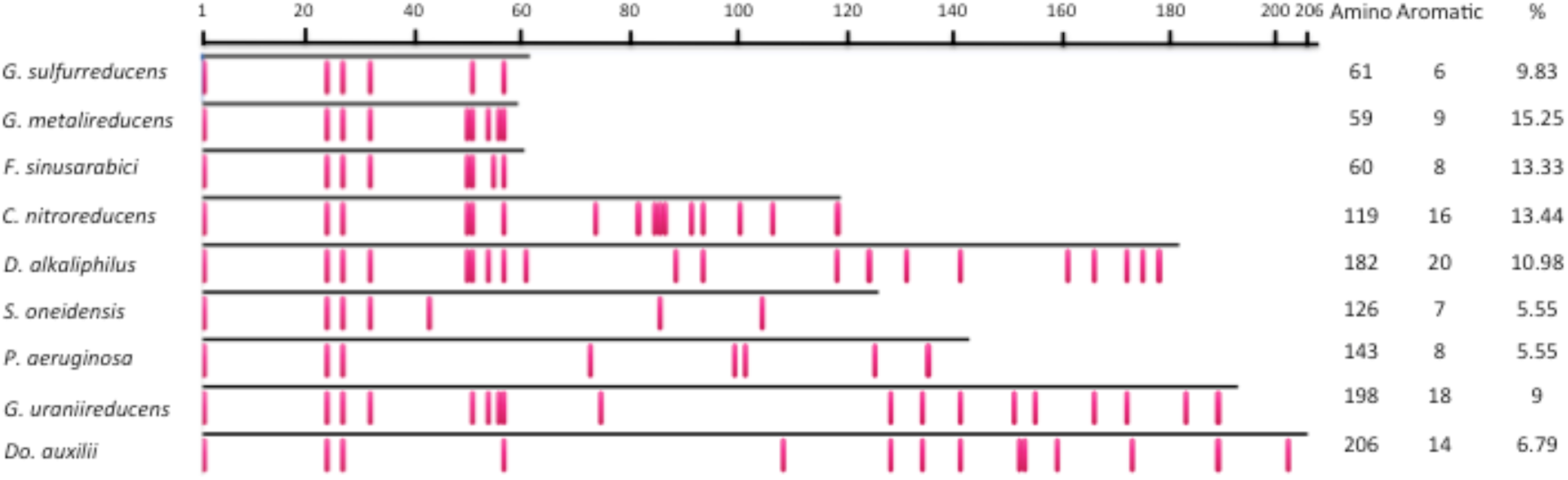
Comparison of aromatic amino acid placement (red rectangles) within mature pilin protein sequences of *Geobacter sulfurreducens*, *Geobacter metallireducens*, *Flexistipes sinusarabici, Calditerrivibrio nitroreducens*, *Desulfurivibrio alkaliphilus*, *Shewcmella oneidensis, Pseudomonas aeruginosa*, *Geobacter uraniireducens*, and *Desulfofervidus auxilii*. Also shown are the total number of amino acids, the number of aromatic amino acids, and the percentage of total amino acids that are aromatic.

However, evaluating pili conductivity is technically challenging, especially with difficult-to-culture microorganisms. For example, it was suggested that e-pili were involved in DIET in a consortium of an ANME-1 microbe and its sulfate-reducing partner (Wegener et al 2015). The sulfate-reducing partner highly expressed a pilin gene when the two microorganisms were growing syntrophically and pili-like filaments were abundant in the co-culture (Wegener et al 2015). It was proposed that the pili of the sulfate reducer were conductive (Wegener et al 2015), but this was not verified, possibly due to difficulties in obtaining sufficient biomass.

The properties of proteins encoded in genomes of difficult-to-culture or as-yet-uncultured microorganisms can often be determined by expressing the genes of interest in more readily cultivated microorganisms. It is possible to screen pilin genes to determine if they have the potential to yield pilin proteins that can assemble into e-pili by heterologously expressing the pilin gene of interest in *G. sulfurreducens* in place of the native *G. sulfurreducens* pilin gene (Liu et al 2014, Tan et al 2016a, Tan et al 2016b, Tan et al 2017, Vargas et al 2013). Only if the pilin gene yields e-pili is *G. sulfurreducens* then capable of producing high current densities on graphite electrodes. For example, pili of *P. aeruginosa* are poorly conductive (Reguera et al 2005) and a strain of *G. sulfurreducens* expressing the pilin gene of *P. aeruginosa* produced low current densities (Liu et al 2014). Expression of the pilin gene of *G. metallireducens* in *G. sulfurreducens* yielded e-pili that were much more conductive than the e-pili originating from the native *G. sulfurreducens* pilin gene (Tan et al 2017). In contrast, the pilin gene of *G. uraniireducens*, which is phylogenetically distinct from the *G. sulfurreducens* and *G. metallireducens* pilin genes (Holmes et al 2016), yielded poorly conductive pili in *G. sulfurreducens* (Tan et al 2016b). As expected, the *G. sulfurreducens* strain expressing the *G. uraniireducens* pilin genes produced low current densities (Tan et al 2016b) as did a *G. sulfurreducens* strain expressing a synthetic pilin gene designed to yield poorly conductive pili (Vargas et al 2013). The poor conductivity of the *G. uraniireducens* pili was consistent with the finding that *G. uraniireducens* employs an electron shuttle for Fe(III) reduction and is incapable of producing high current densities on graphite electrodes (Rotaru et al 2015, Tan et al 2016b).

Therefore, the possibility that diverse microorganisms potentially involved in extracellular electron transfer might have the potential to produce e-pili was evaluated by expressing their pilin genes in *G. sulfurreducens*. The results suggest that pilin gene sequences phylogenetically distinct from the *G. sulfurreducens* pilin gene can yield e-pili.

## Materials and methods

### Bacterial strains, plasmids, and culture conditions

The bacterial strains used in this study are listed in table S1 (Supplementary material). As previously described (Coppi et al 2001), *G. sulfurreducens* was routinely cultured in liquid medium or on agar plates under strict anaerobic conditions in a medium with acetate as the electron donor and fumarate as the electron acceptor at 30 °C. *E. coli* 5-alpha (New England Biolabs, Ipswich, MA, USA) chemically competent cells were used for Gibson Assembly cloning and cultured at 37 °C in Luria-Bertani medium. Gentamicin 20 μg/ml was added to cultures when required for plasmid or chromosomal selection.

### Pilin Gene Sequences and Phylogenetic analyses

Sequence data was acquired from the US Department of Energy Joint Genome Institute (http://www.jgi.doe.gov) or from Genbank at the National Center for Biotechnology Information (NCBI) (http://www.ncbi.nlm.nih.gov). Amino acid alignments were generated with MAFFT (Katoh and Standleyu 2013) and PRANK (Löytynoja and Goldman 2005) algorithms and unreliable residues, columns and sequences in all alignments were identified and eliminated with GUIDANCE2 (Sela et al 2015).

Phylogenetic trees were generated with the maximum likelihood method using MEGA v. 7.0 software (Kumar et al 2016). Before trees were constructed, the Find Best DNA/Protein Models program was run on sequence alignments. The evolutionary history of the PilA protein was inferred with the Maximum Likelihood method based on the Whelan and Goldman model (Whelan and Goldman 2001) and the bootstrap consensus tree was inferred from 100 replicates (Felsenstein 1985). Initial tree(s) for the heuristic search were obtained automatically by applying Neighbor-Joining and BioNJ algorithms to a matrix of pairwise distances estimated using a JTT model, and then selecting the topology with a superior log likelihood value. A discrete Gamma distribution was used to model evolutionary rate differences among sites (5 categories (+G, parameter = 0.9542)). The analysis involved 79 amino acid sequences. All positions with less than 95% site coverage were eliminated. That is, fewer than 5% alignment gaps, missing data, and ambiguous bases were allowed at any position. There were a total of 55 positions in the final dataset.

### Construction of G. sulfurreducens strains expressing heterologous pilins

The strains of *G. sulfurreducens* heterologously expressing the pilin genes of other microorganisms were constructed with a slight modification of the previously described approach (Tan et al 2017). To ensure that the pre-pilin would be correctly recognized and transported to the cellular membrane for processing, the signal peptide from *G. sulfurreducens* (MLQKLRNRKG) was included in all strains. This signal peptide is cleaved during post-translational processing and is not part of the mature pilin sequence. The mature pilin coding sequence from the organisms of interest was inserted downstream from the *G. sulfurreducens* signal peptide sequence in the pYT-1 plasmid (Figure S1). The region between the *SacII* and *NotI* restriction sites, which included the *G. sulfurreducens* promoter region (PpilA), the *G. sulfurreducens* signal peptide sequence, and the heterologous *pilA* sequence, was synthesized as a gene fragment (ThermoFisher Scientific, Waltham MA). Two 15-bp regions homologous to sequences located 5’ and 3’ of the restriction sites on the pYT-1 plasmid were also included in the synthesized gene fragment to facilitate Gibson Assembly. The synthesized DNA was inserted into plasmid pYT-1 between the SacII and NotI restriction sites with the Gibson Assembly Kit ((Gibson et al 2009); New England Biolabs, Ipswich, MA). Correct sequence and assembly was verified by sequencing DNA fragments amplified with *pilA-* Screen-F and pilA-Screen-R primers (Table S2). The recombinant plasmid was then linearized with restriction enzyme SacI and transformed into wild-type *G. sulfurreducens* competent cells by electroporation as previously described (Liu et al 2014).

To ensure that the heterologous gene had correctly integrated into the *G. sulfurreducens* chromosome, sequences were verified with *pilA*-InScreen-F and *pilA*-InScreen-R primers (Table S2). Strain purity was also confirmed by sequencing with this primer set throughout the experiment; i.e. before placing the strain in the bioelectrochemical device, during the current production experiment, and before pili were harvested from cells.

### Current production

The capacity of the strains to produce current was determined as previously described (Nevin et al 2009). Cells were grown in two-chambered H-cell systems with a continuous flow of medium with acetate (10 mM) as the electron donor and graphite stick anodes (65 cm^2^) poised at 300 mV versus Ag/AgCl as the electron acceptor.

### Microscopy

As previously described (Nevin et al 2011), anode biofilms were imaged with confocal laser scanning microscopy using the LIVE/DEAD BacLight viability stain kit (Molecular Probes Eugene, OR, USA). Cells from the anode biofilms were examined by transmission electron microscopy as previously described (Tan et al 2017). Cells placed on carbon-coated copper grids were stained with 2% uranyl acetate and visualized with a JEOL 2000fx transmission electron microscope operated at 200 kV accelerating voltage.

### Pili preparation

Pili were obtained from anode biofilms as previously described (Tan et al 2017). Briefly, pili were sheared from the cells in a blender in 150 mM ethanolamine buffer (pH 10.5), the cells were removed with centrifugation, and the pili were precipitated from the supernatant with the addition of ammonium sulfate, followed by centrifugation. The precipitate was resuspended in ethanolamine buffer, centrifuged to remove additional debris, and the ammonium sulfate precipitation step was repeated. The pili were then resuspended in ethanolamine buffer and stored at 4 °C. Prior to conductance measurements the pili were dialyzed in deionized water at 4 °C with a Slide-A-Lyzer™ MINI Dialysis Device (Thermo Fisher Scientific) with 7 kDa molecular weight cut-off columns. To determine the protein concentration, 20 μl of the pili preparation was air dried, dissociated in 17.2 μl of 1% SDS at pH 1.5 at 95 °C, and then neutralized with 2.8 μl of 1 N NaOH. The protein concentration was analyzed with the Pierce nano bicinchoninic acid assay (BCA assay) (Thermo Fisher, Waltham, MA) All pili preparations were normalized to 500 μg proteins per μl.

### Pili conductance

The method for measuring pili conductance was a modification of the previously described (Malvankar et al 2011, Vargas et al 2013) method for estimating the conductance of pili networks. This method is preferable for screening studies because measuring the conductivity of multiple individual pili is extremely laborious, expensive, and technically difficult. Conductance was measuring with the four-point probe method to eliminate the possibility of contact resistance.

The electrodes were fabricated with photolithography and were 50 nm thick with 10 nm titanium used as an adhesion layer and 40 nm gold (Figure S2). The substrate was 300 nm thick silicon dioxide grown on antimony doped silicon wafers with a wet chemical process. Pads for the four-point probe electrodes were 1 mm by 1mm in length and the electrodes were non-equidistant with 15 μm between the two inner electrodes and 3 μm between each of the inner and outer electrodes. Electrical contacts on the pads were made with 1 μm diameter tungsten probes (Signatone, Gilroy, CA) connected to a Keithley 4200 SCS Parametric Analyzer with triaxial cables.

Pili preparations (2 μl, 500 μg protein/μl) were drop cast onto the gold electrode arrays and dried for 1 h at 24 °C. Then another 2 μl of the pili preparation was drop cast on the electrode array and dried in a desiccator for ca. 12 h.

Each pili preparation was drop cast onto three different devices and current-voltage (I-V) curves for each device were obtained in triplicate with a Keithley 4200 Semiconductor Characterization System (SCS) set up with 4-probes using a +/−30 ×10^−8^ V sweep with a 5 second delay and 250 second hold time. The conductance value of the pili was extracted from the slope of the linear fit of the current-voltage response of the sample as:

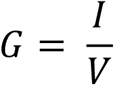

where G is the conductance, I is the current, and V is the voltage.

## Results and discussion

### *e-Pili from the Geobacter-like pilin of* Flexistipes sinusarabici

Many *Geobacter* species and closely related microorganisms have pilin genes that are closely related to the *G. sulfurreducens* pilin gene that yields e-pili (Holmes et al 2016). These include other members of the Desulfuromondales family, such as some *Geoalkalibacter*, *Desulfuromusa,* and *Desulfuromonas* species that are already known to participate in extracellular electron transfer (Holmes et al 2016). However, a few species outside the Desulfuromondales also have pilin genes closely related to those found in *G. sulfurreducens* (Holmes et al 2016). For example, *Flexistipes sinusarabici* is phylogenetically distant from *Geobacter* species based on 16S rRNA gene sequences, but contains a pilin gene closely related to the *G. sulfurreducens* pilin gene (Holmes et al 2016) that is predicted to yield a pilin monomer with a density of aromatic amino acids intermediate between the *G. sulfurreducens* and *G. metallireducens* pilins (Figure 1).

Even under optimal culture conditions *F. sinusarabici* grows poorly, not even producing visible turbidity (Fiala et al 1990). Thus, it was not possible to obtain sufficient biomass to evaluate the conductivity of its pili directly. Therefore, the pilin gene of *F. sinusarabici* was heterologously expressed in *G. sulfurreducens* in place of the native *G. sulfurreducens* pilin gene. The *G. sulfurreducens* strain expressing the *F. sinusarabici* pilin gene grew on anodes, and produced current densities comparable to wild-type *G. sulfurreducens* (Figure 2) with thick biofilms (Figure 3). Such high current densities have only been seen in previous studies when *G. sulfurreducens* is expressing e-pili (Liu et al 2014, Reguera et al 2006, Tan et al 2016b, Tan et al 2017, Vargas et al 2013).

**Figure 2.**
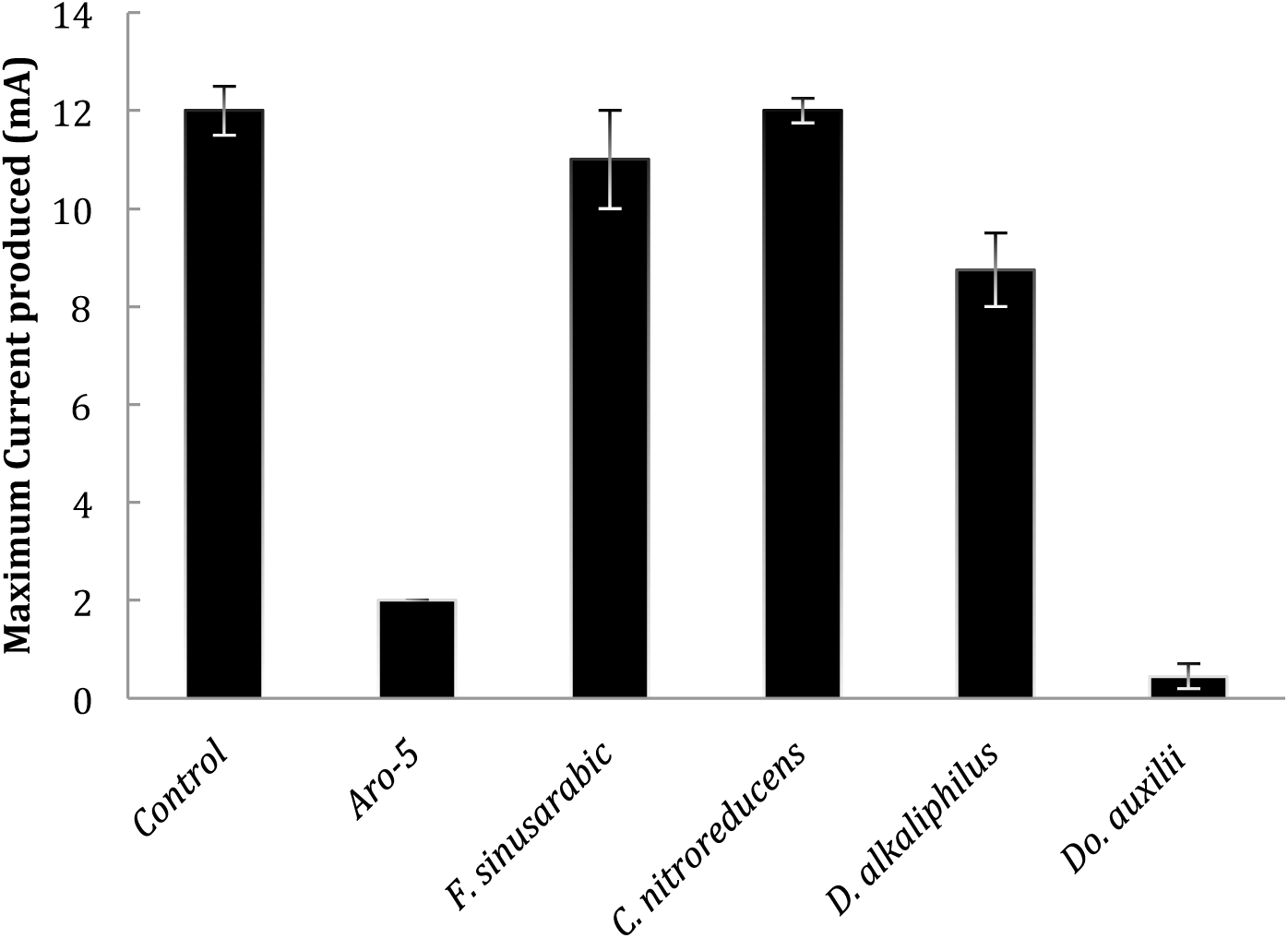
Current production by *G. sulfurreducens* strains expressing either the native wild-type gene (control); a synthetic pilin gene (Aro-5) designed to be poorly conductive (Vargas et al 2013); or the pilin gene of *Flexistipes sinusarabici, Calditerrivibrio nitroreducens*, *Desulfurivibrio alkaliphilus*, or *Desidfofervidus auxilii.* Results shown are the mean of duplicate determinations for each strain with the bars designating the two current values. Aro-5 results from (Vargas et al 2013).

**Figure 3.**
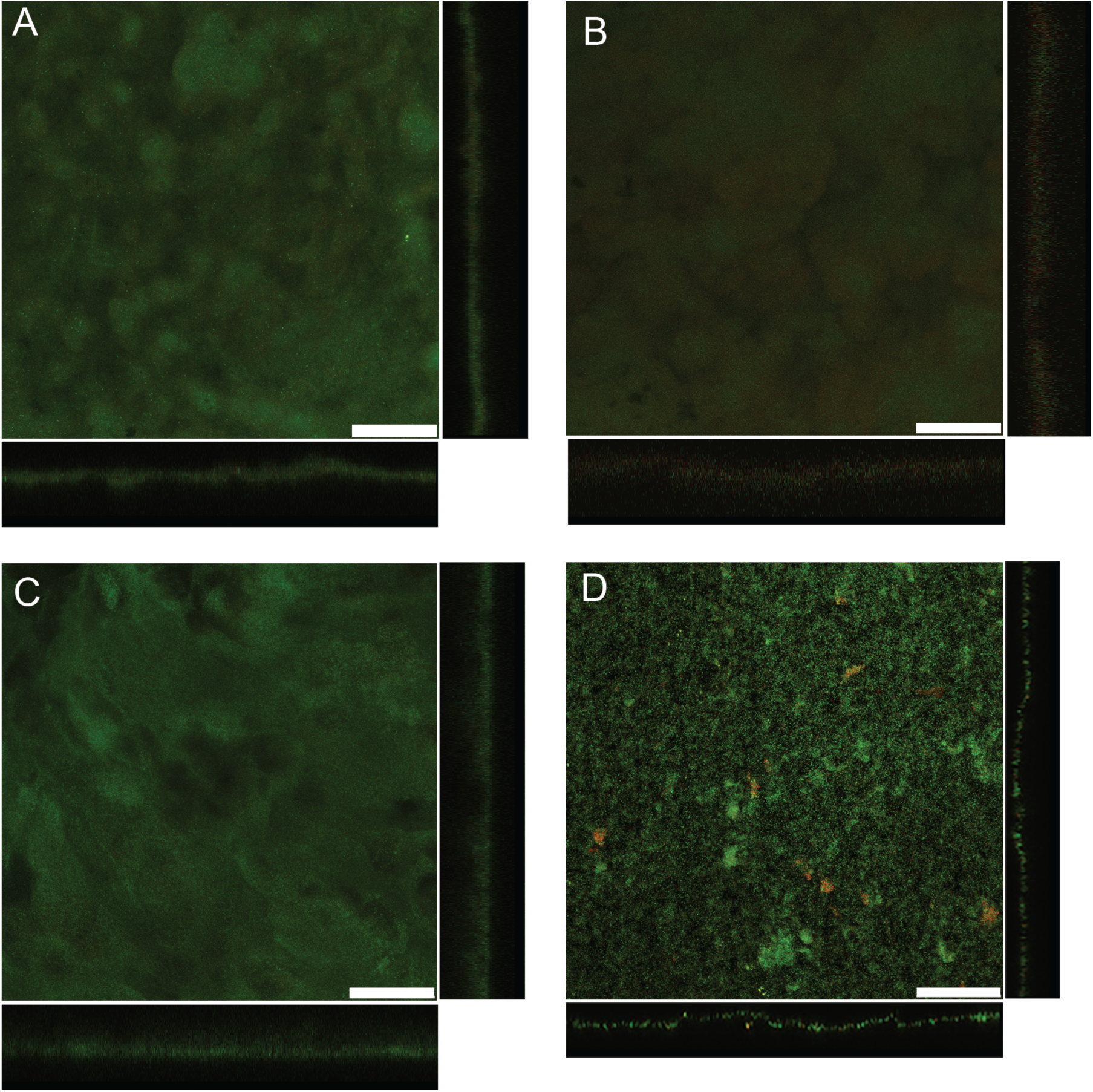
Scanning laser confocal microscopy images of anode biofilms of *G. sulfurreducens* strains heterologously expressing pilin genes from *Flexistipes sinusarabici* (A), *Calditerrivibrio nitroreducens* (B), *Desulfurivibrio alkaliphilus* (C), and *Desulfofervidus auxilii* (D). Top-down three-dimensional, lateral side views (right image) and horizontal side views (bottom image) show cells stained with LIVE/DEAD BacLight viability stain. Bar, 50 μm.

The *G. sulfurreducens* strain expressing the *F. sinusarabici* pilin gene expressed abundant pili (Figure 4A), with a conductance comparable to that of the pili preparations from the control strain of *G. sulfurreducens* expressing its native pilin gene (Figure 5). These results demonstrate that the *F. sinusarabici* pilin gene encodes a pilin monomer that can assemble into e-pili. Close relatives of *F. sinusarabici* include *Geovibrio* and *Deferribacter* species, which are known to be capable of extracellular electron transfer (Alauzet and Jumas-Bilak 2014), and the one available *Deferribacter* genome contains a pilin gene homologous to the *G. sulfurreducens* pilin gene (Holmes et al 2016). Further investigation of the possibility that *F. sinusarabici* employs e-pili for extracellular electron transfer is warranted, but such studies will be challenging because of its poor growth and the lack of tools for genetic manipulation.

**Figure 4.**
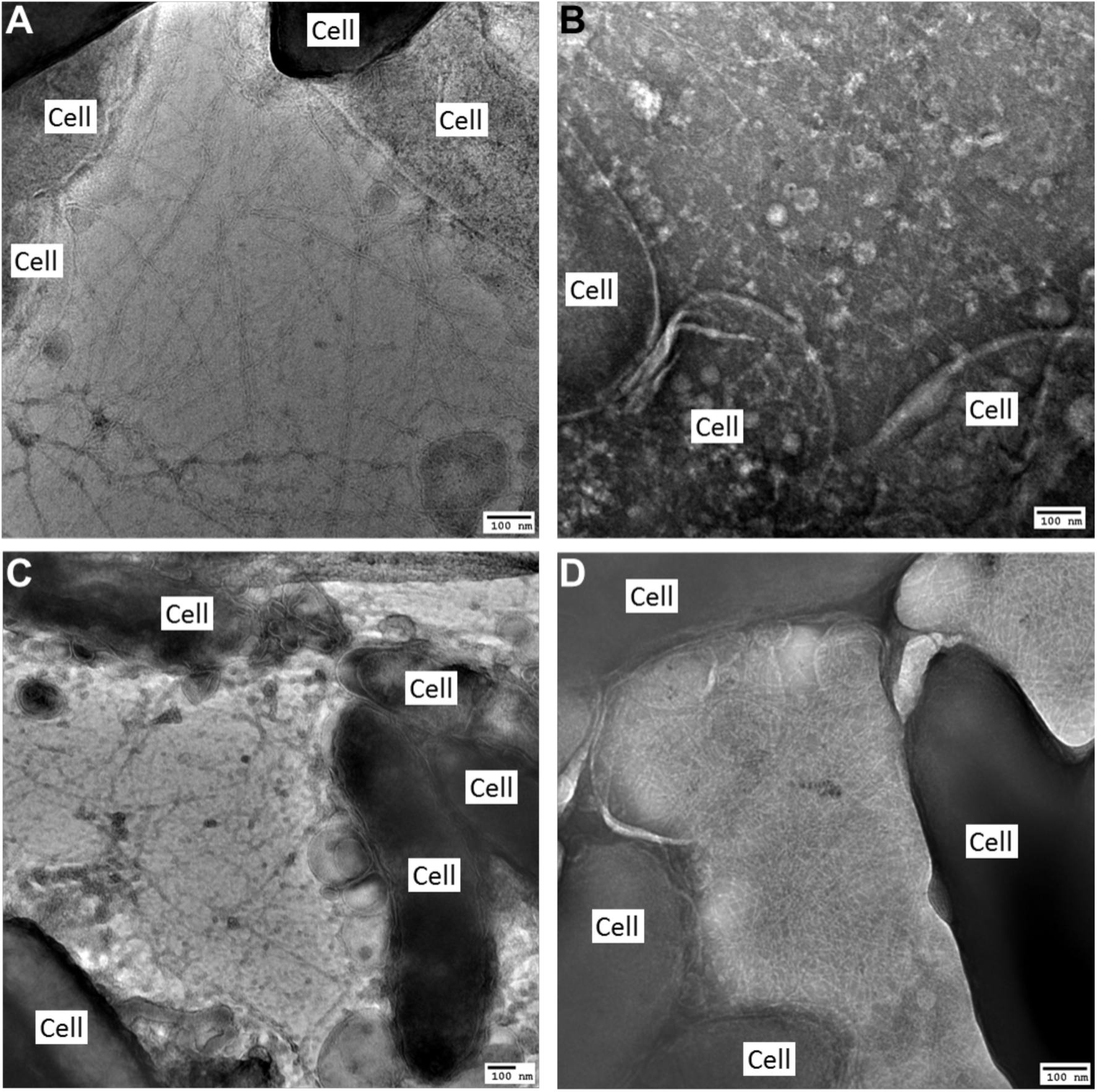
Transmission electron micrographs of *G. sulfurreducens* strains heterologously expressing pilin genes from *Flexistipes sinusarabici* (A), *Calditerrivibrio nitroreducens* (B), *Desulfurivibrio alkaliphilus* (C), and *Desulfofervidus auxilii* (D).

**Figure 5.**
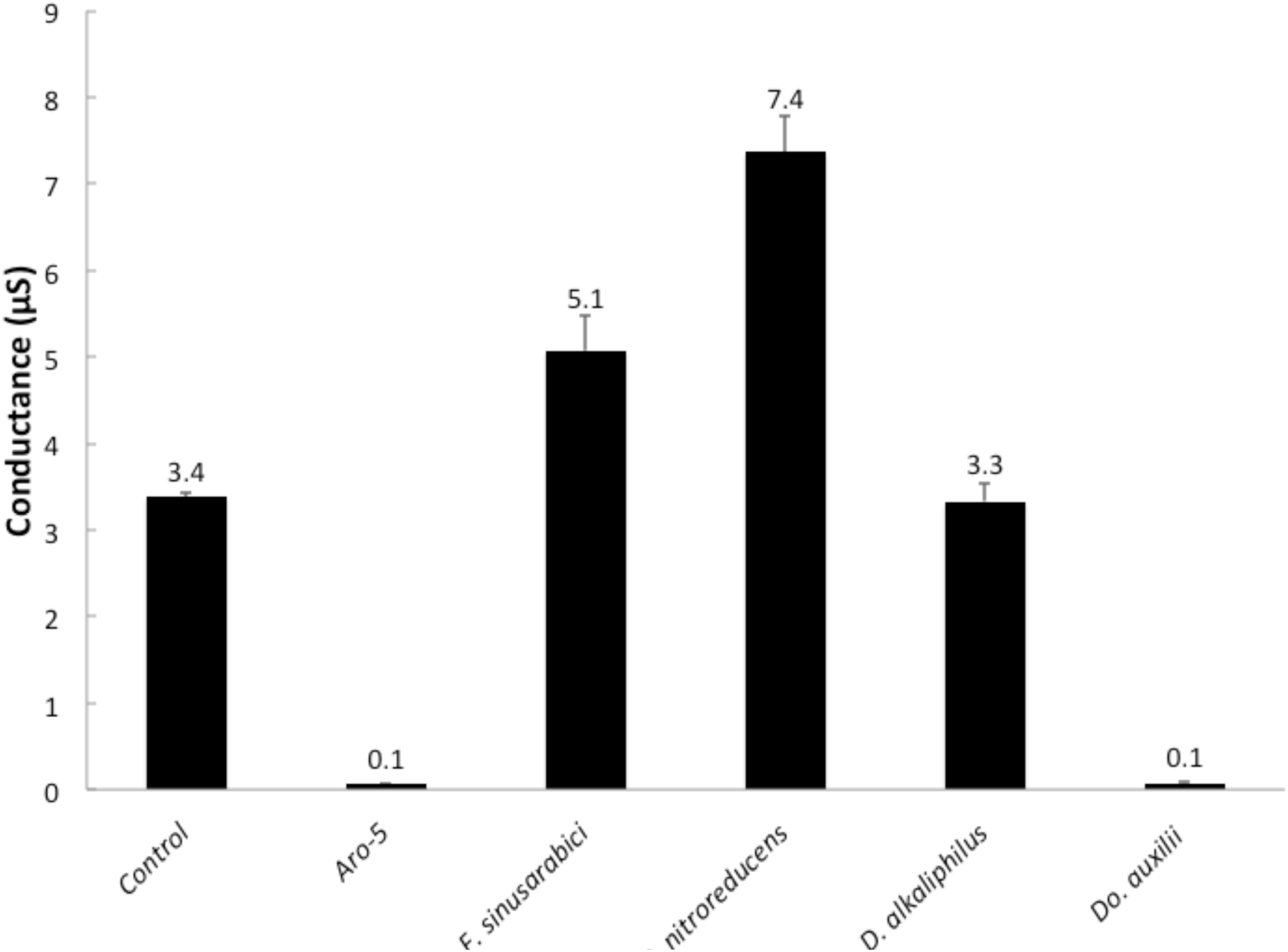
Conductance of pili preparations from *G. sulfurreducens* strains expressing either the native wild-type gene (control); a synthetic pilin gene (Aro-5) designed to be poorly conductive (Vargas et al 2013); or the pilin gene of *Flexistipes sinusarabici, Calditerrivibrio nitroreducens, Desulfurivibrio alkaliphilus,* or *Desulfofervidus auxilii.* Results shown are the mean and standard deviation of triplicate measurements on triplicate electrode arrays.

### e-Pili from phylogenetically distant pilins

As noted in the Introduction, the possibility that pilin genes that are not closely related to the truncated e-pilin gene of *G. sulfurreducens* might also yield e-pili has not been broadly explored. Therefore, this possibility was further investigated with an emphasis on microorganisms that: 1) are thought to be capable of extracellular electron exchange; and 2) have aromatic-rich pilin proteins.

For example, *Calditerrivibrio nitroreducens* is capable of extracellular electron transfer to electrodes (Fu et al 2013) and the density of aromatic amino acids in the *C. nitroreducens* pilin protein is higher than in the *G. sulfurreducens* pilin (Figure 1). However, the *C. nitroreducens* pilin contains nearly twice as many amino acids as the *G. sulfurreducens* pilin. A strain of *G. sulfurreducens* expressing the *C. nitroreducens* pilin gene, rather than the native *G. sulfurreducens* pilin gene, produced current densities similar to the control strain expressing the native pilin gene (Figure 2) with thick biofilms (Figure 3B). The cells produced abundant pili (Figure 4B). The conductance of the pili preparations from the anode biofilm was almost twice that of the control strain expressing the native *G. sulfurreducens* pilin gene (Figure 5). These results demonstrate for the first time that pilin monomers much longer than those of *G. sulfurreducens* can yield e-pili.

Further investigation of the role of e-pili in *C. nitroreducens* will be challenging because of the lack of tools for genetic manipulation of this organism and because this microorganism is difficult to grow to high densities. However, the possibility of *C. nitroreducens* expressing e-pili should be considered when evaluating its ecological niche or possibilities for increasing its current output in microbial fuel cells.

It has been proposed that cable bacteria accept electrons from sulfide-oxidizing bacteria via DIET (Vasquez-Cardenas et al 2015). Genome sequences of cable bacteria have yet to be published but a genome is available for *Desulfurivibrio alkaliphilus* (Melton et al 2016), which is closely related to some species of cable bacteria (Müller et al 2016). The pilin gene of *D. alkaliphilus* is predicted to yield a peptide with even more amino acids than the pilin of *C. nitroreducens* with a density of aromatic amino acids that is comparable to the pilin of *G. sulfurreducens* (Figure 1). Expression of the *D. alkaliphilus* pilin gene in *G. sulfurreducens* yielded a strain that produced thick anode biofilms (Figure 3C) that generated high current densities (Figure 2). Pili were abundant (Figure 4C) with a conductance nearly equal to native *G. sulfurreducens* pili (Figure 5). These results provide another example of a long pilin monomer yielding e-pili.

It does not appear that the ability of *D. alkaliphilus* to reduce Fe(III) oxides or produce current has been examined, but *D. alkaliphilus* does contain porin-cytochrome genes closely related to *Geobacter* genes that are know to be involved in extracelluar electron transfer (Shi et al 2014). Furthermore, *D. alkaliphilus* is capable of growing with elemental sulfur/polysulfide as the electron acceptor (Sorokin et al 2008) and microorganisms capable of elemental sulfur/polysulfide reduction are often capable of electron transfer to Fe(III) or other extracellular electron acceptors (Lovley et al 2004). If cable bacteria closely related to *D. alkaliphilus* have similar pilin genes this might provide a mechanism for the proposed DIET between cable bacteria and sulfur oxidizers (Vasquez-Cardenas et al 2015). The fact that cable bacteria or their putative sulfur-oxidizing partners are not available in pure culture currently limits further investigation into this possibility.

### *Pilin of* Desulfofervidus auxilii *does not yield e-pili*

e-Pili have been proposed to wire an anaerobic methane oxidizer of the ANME-1 phylogenetic group to the sulfate-reducer *Desulfofervidus auxilii* for DIET (Krukenberg et al 2016, Wegener et al 2015). It was not possible for us to obtain a sample of the consortium to directly evaluate the conductivity of the pili, but *G. sulfurreducens* readily expressed pili (Figure 4D) from the *Do. auxilli* pilin gene that was previously reported (Wegener et al 2015) to be highly expressed during syntrophic growth. This strain of *G. sulfurreducens* produced low current densities (Figure 2) with relatively thin biofilms (Figure 3D), and the conductance of the pili was low (Figure 5). There is the possibility that the *Do. auxilii* pilin monomers assemble differently in *Do. auxilii* than they do in *G. sulfurreducens* to yield conductive pili in the natural host. However, this seems unlikely due to the finding that *G. sulfurreducens* expressed e-pili from the pilin monomer of *D. alkaliphilus,* which is of comparable size. Furthermore, the density of aromatic amino acids in the *Do. auxilli* pilin is much lower than that found in e-pili (Figure 1), consistent with the poor conductance of the pili when the *Do. auxilli* pilin gene was expressed in *G. sulfurreducens*. These results suggest that the hypothesis that *Do. auxilii* exchanges electrons with its ANME-1 partner via *Do. auxilii* e-pili (Wegener et al 2015) needs to be revisited. One possible approach would be to measure conductance of the consortium as such measurements can be more technically tractable than studying pili conductance directly (Morita et al 2011, Shrestha et al 2014).

## Conclusions

These results demonstrate that microorganisms outside the *Geobacteraceae* contain pilin genes that can encode e-pili and that pilin monomers much larger than those of *G. sulfurreducens* can yield e-pili. Thus, the diversity of microorganisms that might electrically communicate with other species, or exchange electrons with insoluble minerals, through e-pili may extend well beyond the small percentage of microbes with genes for short e-pilins similar to those found in *G. sulfurreducens*.

The poor understanding of the mechanisms for conductivity for even *G. sulfurreducens* e-pili precludes definitively specifying the features of pilin monomers that confer conductivity to e-pili. However, the limited data set available suggests that pilins with high densities of aromatic amino acids, without large gaps of aromatic-free regions in the pilin monomer, can assemble into e-pili. The phylogenetic distance between the longer pilin sequences found to yield e-pili and the shorter pilin sequences closely related to *G. sulfurreducens* (Figure 6) suggest that aromatic-rich e-pili may have independently evolved several times within bacteria.

**Figure 6.**
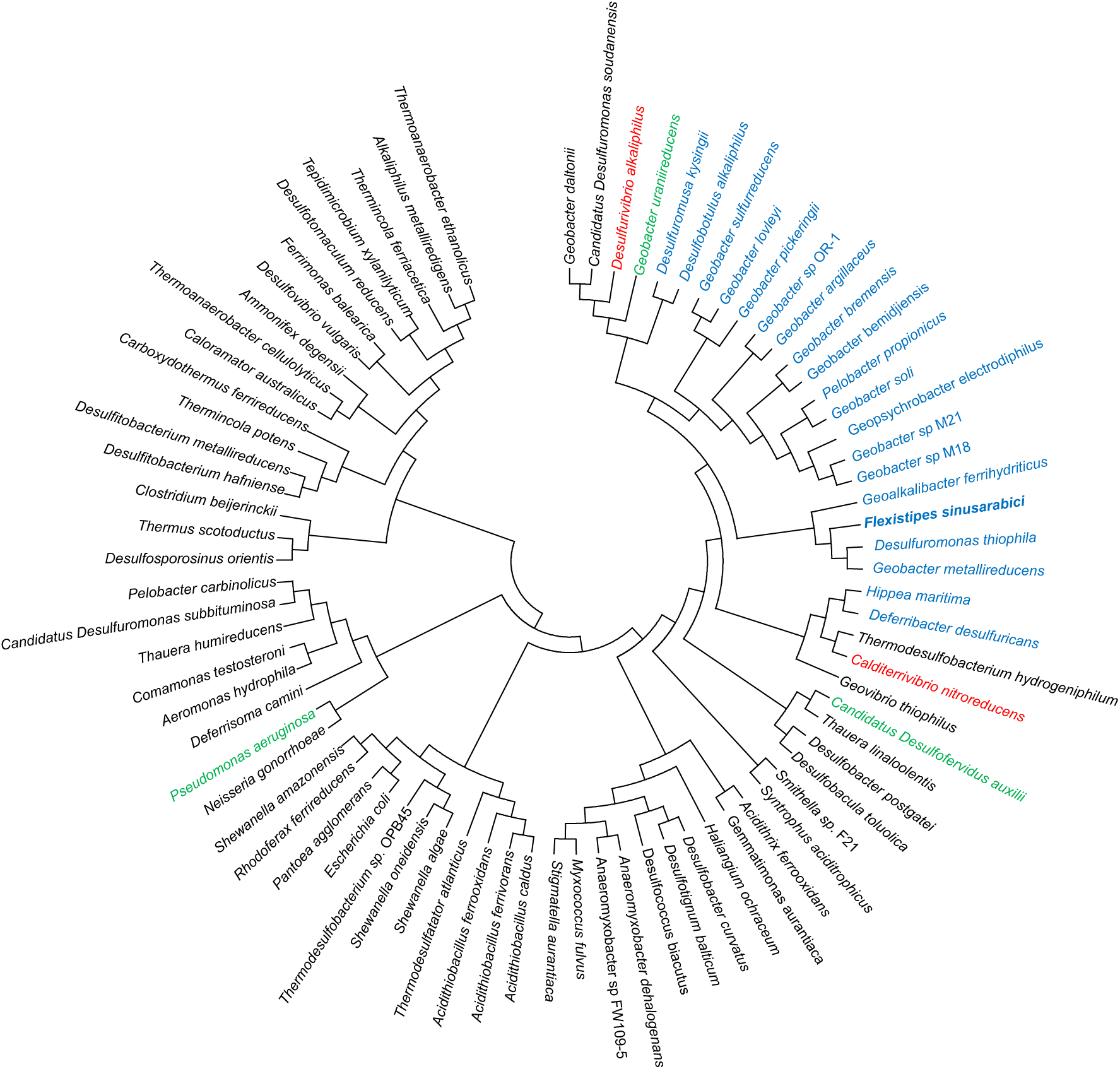
Phylogeny of PilA protein sequences from a diversity of bacteria. Short (ca. 60 amino acids) pilins closely related to *Geobacter sulfitrreducens* are shown in blue, including the pilin from *Flexistipes sinusarabici* (bold). The phylogenetic placement of the longer pilins from *Calditerrivibrio nitroreducens* and *Desulfitrivibrio alkaliphilus* that assembled into conductive pili in *G. sulfitrreducens* are in red. The pilins from *Desulfofervidus auxilii, Shewanella oneidensis,* and *Pseudomonas aeruginosa* that did not yield conductive pili in this or previous studies are shown in green.

The results offer a simple screening strategy to evaluate whether difficult-to-culture microorganisms contain pilin genes that might enable them to participate in extracellular electron exchange with e-pili. The genetic manipulation of *G. sulfurreducens* to heterologously express pilin genes of interest is straightforward and the requirement of *G. sulfurreducens* for e-pili in order to produce high current densities provides a clear phenotype. Thus, the approach described here provides a good first step for hypothesis testing and is preferable to the common practice of suggesting that filaments are “nanowires” based solely on visual appearance.

Diverse microorganisms such as *Aeromonas hydrophila* (Castro et al 2013), *Acidothiobacillus ferroxidans* (Li and Li 2014), *Desulfovibrio desulfuricans* (Eaktasang et al 2016), and *Rhodopseudomonas palustris* (Venkidusamy et al 2015) can produce electrically conductive protein filaments. However, uncertainties about the physiological roles of these filaments exist because: 1) the protein composition of the filaments has not been determined; 2) a role for the filaments in extracellular electron transfer has not been verified; and 3) electron transport along the length of filaments under physiologically relevant conditions has yet to be shown. Expressing pilin genes from these microbes in *G. sulfurreducens* may help better define these filaments.

e-Pili are a potential source of ‘green’ electronic materials that can be sustainably mass-produced with renewable feedstocks without toxic components in the final product (Adhikari et al 2016, Tan et al 2016a, Tan et al 2017). The e-pili described here greatly expand the options for starting materials and the screening method described will facilitate searching the microbial world, including the metagenomes of uncultured microorganisms, for additional conductive materials.

## Acknowledgements

This research was supported by Office of Naval Research Grants N000141310549 and N000141612526.

## Supplementary Material

**Walker et al. Electrically Conductive Pili from Pilin Genes of Phylogenetically Diverse Microorganisms**

**Table S1.** The bacterial strains used in this study.

| Strain or plasmid | Relevant characteristic(s) | Source or reference |
| --- | --- | --- |
| **<u>Strains</u>** |  |  |
| ***E. coli*** |  |  |
| NEB 5-alpha E. coli | *fhuA2 Δ(argF-lacZ)U169 phoA glnV44 Φ80 Δ(lacZ)M15 gyrA96 recA1 relA1 endA1 thi-1 hsdR17* | NEB, Ipswich, MA |
| ***G. sulfurreducens*** |  |  |
| PCA | Wild type | Lab stock ENREF_1 |
| PCA-Gen | PCA control strain expressing *Gm<sup>r</sup>* and wild type *pilA* | Lab stock ENREF_1 |
| PCA-FS | PCA expressing *F. sinusarabici pilA* allele; *Gm<sup>r</sup>* | This work |
| PCA-CN | PCA expressing *C. nitroreducens pilA* allele; *Gm<sup>r</sup>* | This work |
| PCA-DA | PCA expressing *D. alkaliphilus pilA* allele; *Gm<sup>r</sup>* | This work |
| PCA-DoA | PCA expressing *Do. auxilii pilA* allele; *Gm<sup>r</sup>* | This work |
| **<u>Plasmids</u>** |  |  |
| pYT-1 | Upstream and downstream homology arms containing PCA-<br>*pilA* and GSU1497 coding sequences; *gm<sup>r</sup>loxP*, *Gm<sup>r</sup>*, *Cm<sup>r</sup>* | Tan *et al* (2017) |
| pYT -FS | pYT-1 carrying the *PpilA*, *F. sinusarabici pilA* allele and<br>GSU1497 coding sequence; *Gm<sup>r</sup>*, *Cm<sup>r</sup>* | This work |
| pYT -CN | pYT-1 carrying the *PpilA*, *C. nitroreducens pilA* allele and<br>GSU1497 coding sequence; *Gm<sup>r</sup>*, *Cm<sup>r</sup>* | This work |
| pYT -DA | pYT-1 carrying the *PpilA*, *D. alkaliphilus pilA* allele and<br>GSU1497 coding sequence; *Gm<sup>r</sup>*, *Cm<sup>r</sup>* | This work |
| pYT -DoA | pYT-1 carrying the *PpilA*, *Do. auxilii pilA* allele and GSU1497<br>coding sequence; *Gm<sup>r</sup>*, *Cm<sup>r</sup>* | This work |

*Gm<sup>r</sup>*, Gentamicin resistance, *Cm<sup>r</sup>*, Chlorophenol resistance .

**Table S2.**
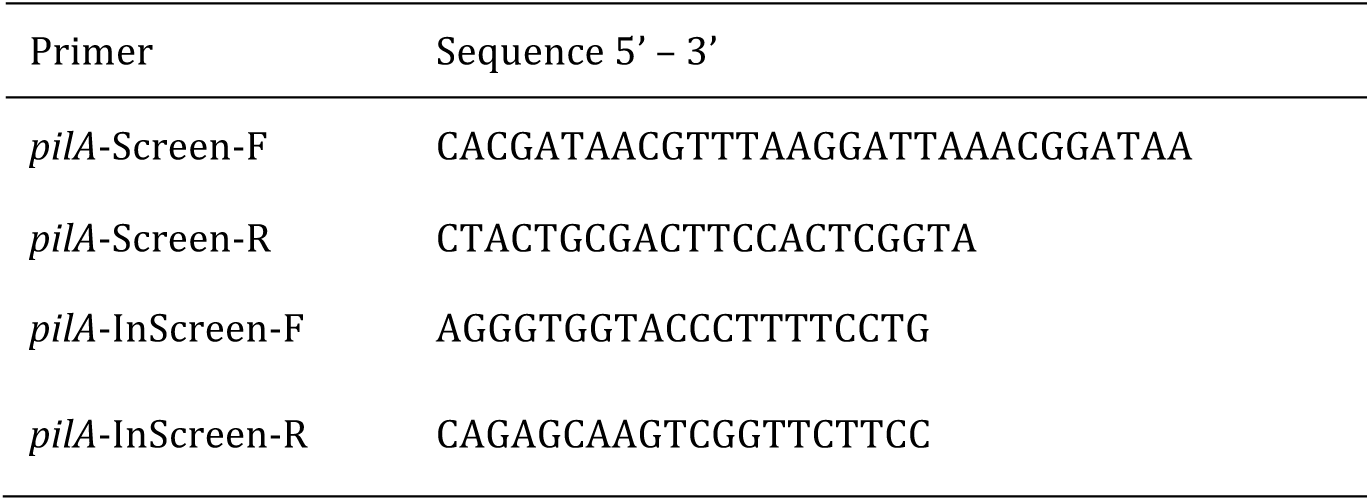
The bacterial strains used in this study.

**Supplementary Figure S1.**
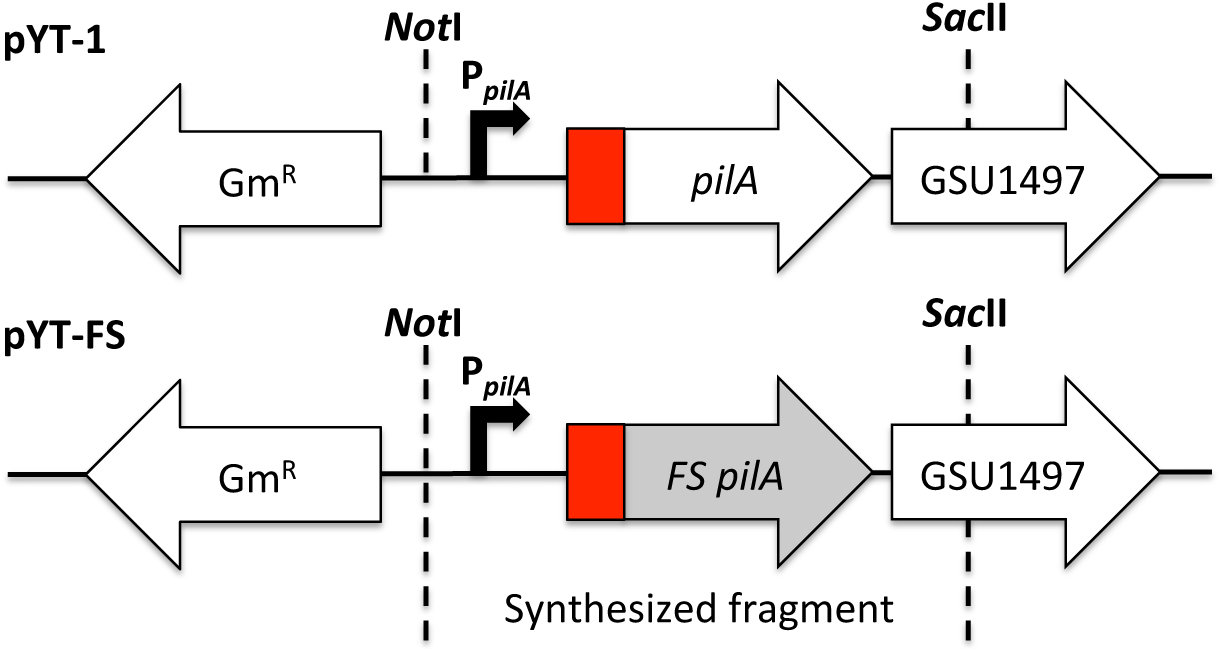
Region on the plasmid pYT-1 which was synthesized and cloned between the *Not*I and *Sac*II restriction sites. Included in the synthesis are the P*_pilA_,* the native signal peptide for the *pilA* (red), and the heterologous mature pilA encoding sequence (grey FS-pilA), as well as part of GSU1497 downstream homology arm.

**Supplementary Figure S2.**
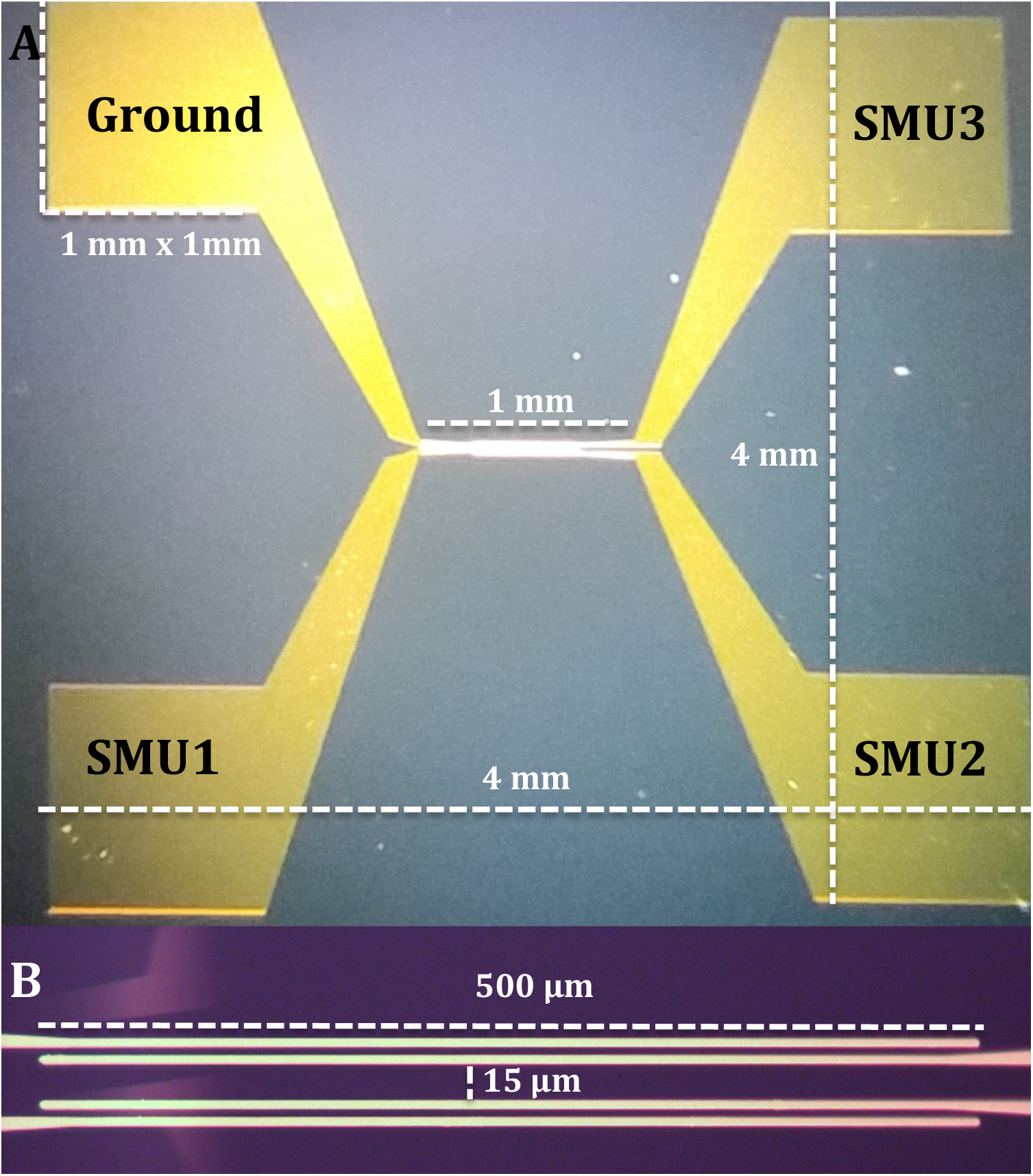
Diagram showing the architecture of the electrode devices used. There is a 300 nm oxide layer with 40 nm gold electrodes on 10 nm thickness tungsten. Current is applied at SMU1 and is removed at Ground. The voltage difference is measured between SMU2 and SMU3. A) The diameter of the device is 4 mm with each gold pad measuring 1 mm × 1mm. B) Close up of the 4 electrodes. The distance between SMU1 - SMU2 and SMU3 – Ground is 3 μm, and the distance between SMU2 and SMU3 is 15 μm.

Vargas, M., Malvankar, N. S., Tremblay, P. L., Leang, C., Smith, J. A., Patel, P., et al. (2013). Aromatic amino acids required for pili conductivity and long-range extracellular electron transport in *Geobacter sulfurreducens.* mBio 4:e00105-13. doi:10.1128/mBio.00105-13

Tan Y, Adhikari RY, Malvankar NS, Ward JE, Woodard TL, Nevin KP, Lovley DR. (2017). Expressing the Geobacter metallireducens PilA in Geobacter sulfurreducens yields pili with exceptional conductivity. mBio 8:e02203-16. https://doi.org/10.1128/mBio.02203-16.

